# Divergent gene expression responses in two Baltic Sea heterotrophic model bacteria to dinoflagellate dissolved organic matter

**DOI:** 10.1101/2020.11.23.393876

**Authors:** Christofer M.G. Osbeck, Daniel Lundin, Camilla Karlsson, Jonna E. Teikari, Mary Ann Moran, Jarone Pinhassi

**Affiliations:** Centre for Ecology and Evolution in Microbial Model Systems, EEMiS, Linnaeus University, Kalmar, Sweden; Department of Microbiology, University of Helsinki, Helsinki, Finland; Department of Marine Sciences, University of Georgia, Athens, Georgia, USA

**Keywords:** Marine bacteria, bacterioplankton, microbial ecology, genomics, transcriptomics, mRNA, RNAseq, phytoplankton exudation, ecological traits

## Abstract

Phytoplankton release massive amounts of dissolved organic matter (DOM) into the water column during recurring blooms in coastal waters and inland seas. The released DOM encompasses a complex mixture of both known and unknown compounds, and is a rich nutrient source for heterotrophic bacteria. The metabolic activity of bacteria during and after phytoplankton blooms can hence be expected to reflect the characteristics of the released DOM. We therefore investigated if bacterioplankton could be used as “living sensors” of phytoplankton DOM quantity and/or quality, by applying gene expression analyses to identify bacterial metabolisms induced by DOM. We used transcriptional analysis of two Baltic Sea bacterial isolates (*Polaribacter* sp. BAL334 [Flavobacteriia] and *Brevundimonas* sp. BAL450 [Alphaproteobacteria]) growing with DOM from axenic cultures of the dinoflagellate *Prorocentrum minimum*. We observed pronounced differences between the two bacteria both in bacterial growth and the expressed metabolic pathways in cultures exposed to dinoflagellate DOM compared with controls. Differences in metabolic responses between the two isolates were caused both by differences in gene repertoire between them (e.g. in the SEED categories for membrane transport, motility and photoheterotrophy) and the regulation of expression (e.g. fatty acid metabolism), emphasizing the importance of separating the responses of different taxa in analyses of community sequence data. Similarities between the bacteria included substantially increased expression of genes for Ton and Tol transport systems in both isolates, which are commonly associated with uptake of complex organic molecules. *Polaribacter* sp. BAL334 showed stronger metabolic responses to DOM harvested from exponential than stationary phase dinoflagellates (128 compared to 26 differentially expressed genes), whereas *Brevundimonas* sp. BAL450 responded more to the DOM from stationary than exponential phase dinoflagellates (33 compared to 6 differentially expressed genes). These findings suggest that shifts in bacterial metabolisms during different phases of phytoplankton blooms can be detected in individual bacterial species and can provide insights into their involvement in DOM transformations.

## Introduction

Dissolved organic matter (DOM) in seawater is estimated to represent one of the largest reservoirs of organic carbon on earth (1). It consists of a complex mixture of compounds of different molecular weights, solubility and volatility (2) and is traditionally classified according to bioavailability (i.e. labile, semi-labile and refractory) with turnover times ranging from minutes to thousands of years (3). DOM released by living and dying phytoplankton is an important source of organic carbon available for heterotrophic bacteria (4). Up to 50% of the carbon fixed by primary producers in marine and limnic ecosystems – bacterial and eukaryotic phytoplankton as well as multicellular algae – is turned over by bacterioplankton in the microbial loop (5). This way, organic carbon is degraded and transformed by the microbial community, with most eventually respired as CO_2_. This carbon turnover occurs at a rate that is orders of magnitude higher in the sea than in terrestrial ecosystems (6, 7), particularly so in coastal environments and inland seas where nutrient concentrations do not limit microbial activities to the same extent as in the open ocean. Given the tight linkages between phytoplankton and bacteria, it is desirable to learn to what extent and by which mechanisms the metabolic activity of heterotrophic bacteria regulate carbon and nutrient cycling through the microbial loop.

Monitoring the actual rates of the plethora of metabolic pathways active in microbial communities directly *in situ* is currently not feasible, but nucleotide sequencing-based methods, in particular metatranscriptomics, can indicate which microbial metabolisms are actively transcribed. Metatranscriptomics has hence become a widely used tool to provide detailed insights into the genetic underpinnings of metabolic responses within communities both in natural environments and controlled experiments (8–11). The complexity of gene regulation observed in communities consisting of many thousands of individual populations is, however, daunting. For the purpose of eventually using transcript profiling as a proxy for metabolic activity in complex natural communities, zooming in to compare gene expression pattern in isolates of environmentally relevant microbial taxa could be useful.

With the aim of charting the possibility of using transcriptional activity of bacterial isolates as living sensors for the flow of nutrients in the ecosystem, we exposed two Baltic Sea model bacteria to DOM derived from axenic cultures of *Prorocentrum minimum*, a dinoflagellate that forms major blooms in the spring and autumn in the Baltic Sea. The bacterial isolates – *Polaribacter* sp. strain BAL334 (*Flavobacteriaceae, Bacteroidetes*) and *Brevundimonas* sp. strain BAL450 (*Caulobacteraceae*, *Alphaproteobacteria*) – were selected to compare responses of bacteria with different evolutionary histories. Furthermore, since previous studies of phytoplankton extracellular DOM release have suggested that phytoplankton secrete different compounds during early and late growth phases (12–16), the DOM was harvested both from dinoflagellates growing actively and in stationary phase. This allowed characterization of potential differences in bacterial responses to DOM released during exponential and senescence phases of phytoplankton blooms.

## Materials and Methods

### Cultivation of axenic Prorocentrum minimum

An axenic culture of the dinoflagellate *Prorocentrum minimum* strain CCMP1329 was obtained from the Provasoli-Guillard National Center of Marine Algae and Microbiota (CCMP; https://ncma.bigelow.org/). 5 mL of inoculum of *P. minimum* CCMP1329 was transferred and cultivated in axenic conditions in 6 replicates using acid-washed Erlenmeyer flasks (2 L) containing 1.3 L of L1 medium (17), prepared using 0.2 μm membrane filters (Supor®, Pall Corporation) and artificial seawater (30 practical salinity units, prepared from Sea Salts; Sigma). The cultures were placed in 20°C with photosynthetically active radiation (PAR) of 83 – 101 μmol photon m^−2^ s^−1^ in light:dark cycles of 13:11 h and bubbled with filtered air provided by an inhouse air system. To follow the growth of the cultures, chlorophyll *a* concentrations were measured regularly by collecting 1 mL of culture on 25 mm glass microfiber filters (GF/C, Glass Microfiber Binder Free, Whatman), followed by chlorophyll *a* ethanol extraction according to (18).

### Collection of DOM

DOM from *P. minimum* CCMP1329 was collected from three of the cultures in the exponential growth phase (~15 days after inoculation) and from two of the cultures in stationary phase (~31 days), hereafter referred to as DOM_exp and DOM_sta, respectively. DOM from the exponential growth phase was retrieved as follows. Phytoplankton cells were gently removed by first filtering through an acid-washed 3.0 μm polycarbonate filter (GSV, Life Science) and then through an acid-washed 0.22 μm polycarbonate filter (GSV, Life Science), using a Sterifil 47 mm filter holder (Merck Millipore). DOM collected in stationary growth phase was obtained by first centrifuging the cultures in acid-washed 50 mL Falcon tubes for 10 min at 3000 g (to prevent filters from clogging); the supernatant was then filtered through an acid-washed 0.22 μm polycarbonate filter (GSV, Life Science) using a Sterifil 47 mm filter holder (Merck Millipore). The flow-through liquid was transferred into an acid-washed 10 L polycarbonate (PC) bottle and mixed before samples for dissolved organic carbon (DOC) concentration and microscope samples were taken (see below for detailed information of the sampling procedure). Finally, the DOM was aliquoted into 1 L acid-washed PC bottles and stored at −80°C until further proceedings.

### Bacterial isolates and culture conditions

The flavobacterium *Polaribacter* sp. strain BAL334 (hereafter referred to as *Polaribacter* BAL334) and alphaproteobacterium *Brevundimonas* sp. strain BAL450 (hereafter referred to as *Brevundimonas* BAL450) were isolated from surface water (2 m depth) at the Linnaeus Microbial Observatory (LMO) in the Baltic Sea (N 56° 55.8540', E 17° 3.6420') during 2012. Seawater was spread on Baltic Zobell agar plates containing a mixture of 5 g bacto peptone, 1 g yeast extract and 15 g bacto agar per L of sterile Baltic Sea water (i.e. a mix of 750 ml seawater and 250 ml MilliQ water). Bacterial colonies were transferred into 1 mL of Baltic Zobell medium (i.e. mixture of 5 g bactopeptone and 1 g yeast extract per L of sterile Baltic Seawater) and preserved in glycerol (25%, final concentration) in −80°C.

### DNA extraction and genome sequencing

To identify the bacteria, DNA from the isolates were extracted using the E.Z.N.A. Tissue DNA kit (Omega bio-tek, USA) following the manufacturer's protocol for extraction of cultured cells in suspension. For identification of the isolates, bacterial 16S rRNA genes were PCR amplified using the primers 27F and 1492R at a final concentration of 10 picomole per μl with the following PCR thermal cycling program: 95°C for 2 min; 30 cycles of 95°C for 30 s, 50°C for 30 s, and 72°C for 45 s; and 72°C for 7 min. E.Z.N.A. Cycle-Pure Kit (Omega bio-tek, USA) were used for cleaning the PCR product following the manufacturer's spin protocol instruction. Samples were sent for Sanger sequencing at Macrogen Europe, Amsterdam, Netherlands. The partial 16S rRNA gene sequences have been deposited in GenBank with the following accession numbers: KM586879 (*Polaribacter* BAL334) and KM586934 (*Brevundimonas* BAL450).

Genome sequences from the isolates *Polaribacter* BAL334 and *Brevundimonas* BAL450 were obtained by sequencing the extracted genomic DNA using the Illumina HiSeq 2500 system (PE 2×125bp) at SciLifeLab, Solna, Sweden. The quality of sequences was checked with FastQC (version 0.11.5) (19) and MultiQC (version 1.4) (20), adaptors were removed with cutadapt (version 1.12) (21) and Sickle (version 1.33) (22) was used to trim sequences based on quality score. Assembly was performed using Megahit (version 1.1.2) (23) and annotation with the Rapid Annotation using Subsystem Technology (RAST) server (24, 25). The genomes are available in the RAST database SEED viewer (https://rast.nmpdr.org/seedviewer.cgi) with identities 6666666.325781 (https://rast.nmpdr.org/seedviewer.cgi?page=Organism&organism=6666666.325781) and 6666666.325780 (https://rast.nmpdr.org/seedviewer.cgi?page=Organism&organism=6666666.325780) for *Polaribacter* BAL334 and *Brevundimonas* BAL450, respectively, using the guest account (user login “guest”, password “guest”).

### Experimental setup and spiking of DOM

Bacterial isolates were grown on Baltic Zobell agar plates for 3-4 days at room temperature after being transferred from the −80°C freezer. Subsequently, they were inoculated into an acid-washed 100-mL glass bottle containing 20 mL Baltic Zobell medium, and were allowed to grow for 28 hours (*Polaribacter* BAL334) and 54 hours (*Brevundimonas* BAL450) on a Unimax 2010 orbital shaker (Heidolph) at 120 rpm. 2 mL of bacterial culture *Polaribacter* BAL334 (reaching an optical density [OD_600_] of 0.39) and 0.5 mL of bacterial culture *Brevundimonas* BAL450 (reaching OD_600_ of 0.57) were transferred to acid-washed 2 (L) glass bottles containing 300 mL fresh Baltic Zobell medium. *Polaribacter* BAL334 was grown into early stationary phase at 80 rpm (38 hours; and OD_600_ 0.20) and *Brevundimonas* BAL450 bacterial cultures was grown into early stationary phase (40 hours; OD_600_ 1.3) at 120 rpm. To reduce nutrient concentrations and adapt bacterial cells to starvation, bacterial cells were centrifuged at 4000 rpm for 7 min, supernatants were discarded and cell pellets were washed by adding 1 volume of sterile artificial Baltic seawater (7 PSU, prepared from Sea Salts; Sigma). Cells were resuspended in artificial Baltic seawater and the procedure was repeated twice more.

To start the experiment, 30 mL (*Polaribacter* BAL334) and 14.5 mL (*Brevundimonas* BAL450) of resuspended bacterial cells were divided into each of nine acid-washed (1 L) glass bottles containing 700 mL artificial seawater (7 practical salinity units, prepared from Sea Salts; Sigma). The different volumes were to start the experiment with a similar bacterial biomass, and was based on OD measures in the washed cells (see previous paragraph). Subsequently, after 1.5 hours, three of the bottles were spiked with 67 mL DOM_exp and three with 14 mL DOM_sta to obtain an enrichment with DOC corresponding to ~50 μM carbon (final concentration). To minimize potential effects from the medium used for culturing *P. minimum* between the treatments, 53 mL of L1 medium were added to the DOM_sta bottles. Three bottles serving as controls were spiked with 67 mL of L1 medium. Samples for determination of DOC concentrations and bacterial abundances were taken as described below.

### Determination of DOC concentrations, bacterial abundance, OD and purity of cultures using microscopy and cultivation

Samples for DOC concentrations were collected 1 hour after DOM spiking by filtering 30 mL of sample through 0.2 μm acrodisc supor syringe filters 32 mm, into a 60 mL TC flask (Sarstedt) using a 50 ml plastic syringe (NORM-JECT). Samples were then acidified by addition of 448 μl 1.2 M HCl. DOC concentrations were calculated as non purgeable organic carbon (using high temperature catalytic oxidation followed by NDIR detection of the gaseous CO_2_), analyzed on the high-temperature carbon analyzer Shimadzu TOC-5000 at Umeå Marine Science Centre, Umeå, Sweden. Bacterial abundance (BA) samples were taken in triplicates from each replicate 1 hour after DOM spiking, by fixing the sample with paraformaldehyde at a final concentration of 1%. Samples were then frozen at −80°C until determined by using the flow cytometer Cube 8 (CyFlow®) according to protocol in (26). Optical density at 600 nanometer (OD_600_) was measured with a Beckman DU®640 spectrophotometer. To ensure axenic conditions (i.e. exclusion of bacterial contamination), 1 mL of algae cultures and samples from experiments were fixed with 1% paraformaldehyde, stained with 0.02% SYBR gold (final concentration) and filtered onto 0.2 μm, 25 mm black polycarbonate filters (Millipore). The filters were analyzed by epifluorescence microscope observation before and after the cultivations/experiments. Additionally, aliquots from bacterial and phytoplankton cultures were spread on Zobell agar plates for investigation of potential contamination.

### RNA sampling, extraction and sequencing

One hour after DOM spiking, seawater samples for RNA were fixed by addition of an ethanol:phenol mix (5 % phenol in absolute ethanol) in a 10:1 proportion (27). The fixed samples were then filtered through Durapore 0.2 μm, 47-mm membrane filters GV (Merck Millipore). Filters were folded and transferred into clean nucleotide-free collection tubes and stored at −80°C until further procedure. Extraction of RNA was performed using the RNeasy mini kit (Qiagen). Briefly, bacterial cells were lysed by cutting membrane filters into smaller pieces and placing them in nucleic acid free microfuge tubes containing RLT Buffer (with added ß-Mercaptoethanol 1:100) and 1.5 gram 200 μm Low Binding Zirconium Beads (OPS diagnostics, USA). Thereafter, cell lysis, RNA purification, on column DNase digestion and RNA elution were performed following the manufacturer's instructions. Total RNA was DNase treated using the TURBO DNA-free Kit (ThermoFisher Scientific) and quality checked on agarose gel. Ribosomal RNA was depleted using RiboMinus Transcriptome Isolation Kit and RiboMinus Concentration Module (ThermoFisher Scientific) and mRNA was amplified using the MeassageAmp II-Bacteria RNA Amplification Kit (ThermoFisher Scientific) according to manufacturer's instructions. RNA sequencing was done at the at SciLifeLab, Solna, Sweden. Raw sequence reads are available at NCBI’s Sequence Read Archive under the BioProject PRJNA678611 (https://www.ncbi.nlm.nih.gov/bioproject/PRJNA678611).

### Bioinformatics and statistical analysis

RNA sequencing was done at the at SciLifeLab, Solna, Sweden, and RNA sequence reads were quality trimmed with Sickle (version 1.33) (22) and mapped to the genomes with Bowtie 2 (28). This resulted in between 92,655 and 329,457 mRNA sequence reads per sample. EdgeR (29) was used to determine significantly differentially expressed genes (false discovery rate <5%) between treatments and controls. EdgeR was also used to retrieve normalized counts per million (CPM) estimates. Genes with an expression of less than five sequence reads were not included in the analyses.

## Results

### Growth of Prorocentrum minimum

The dinoflagellate *Prorocentrum minimum* was grown as the DOM source for the experiments. It followed a sigmoid growth curve with a lag phase of about 10 d. Thereafter, it entered an exponential growth phase. DOM was collected during active growth (day 15) at a chlorophyll *a* concentration of 629 ± 35 μg/L. After 31 d the samples for stationary phase DOM were retrieved at chl *a* concentration of 2436 ± 130 μg/L (**Fig. 1**).

**Figure 1.**
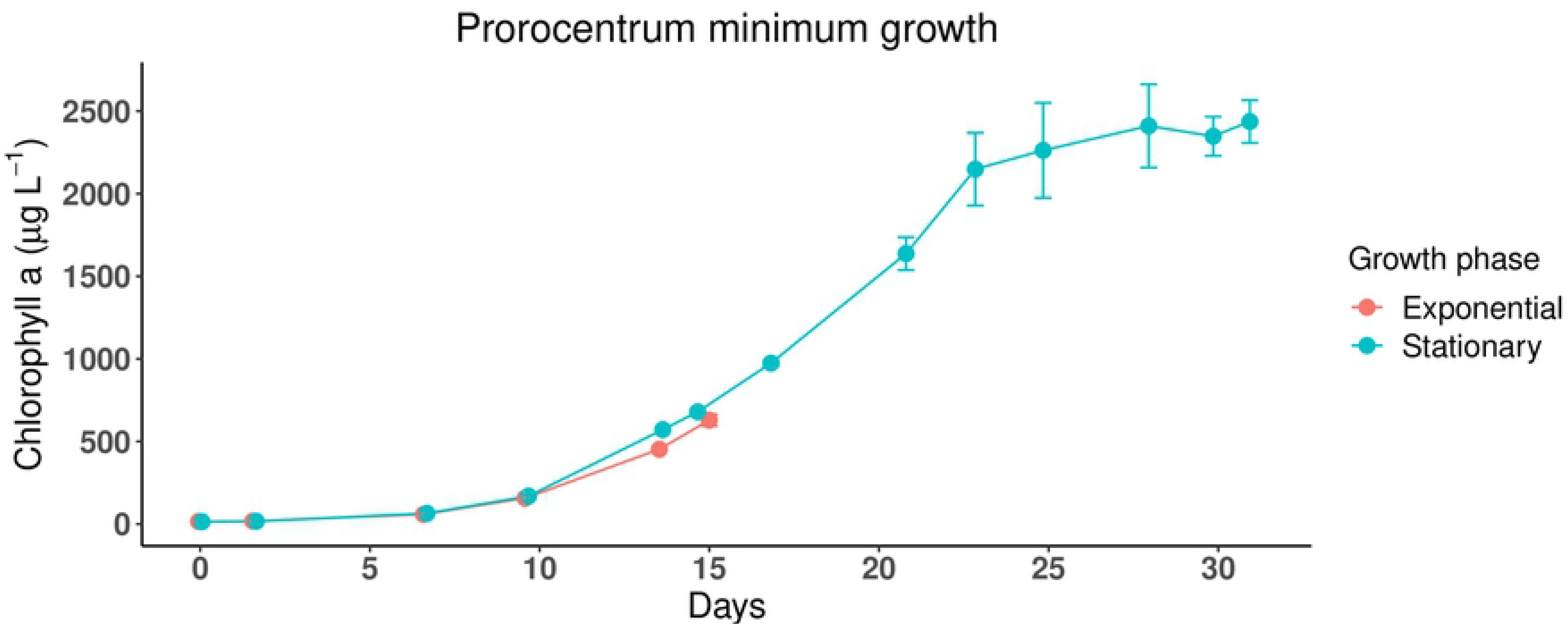
Growth of axenic *P. minimum* cultures in L1 medium for collection of dissolved organic matter from different growth phases. Red line denotes growth of cultures harvested for DOM in exponential phase, and blue line shows growth of cultures harvested in stationary phase. Chlorophyll *a* concentrations were monitored as a proxy for biomass. Error bars denote the standard deviations of triplicates for exponential phase cultures and duplicates for stationary phase; when not visible, error bars are within symbols.

### Bacterial abundance

In *Polaribacter* BAL334 cultures spiked with DOM from exponential and stationary phase dinoflagellate cultures, bacterial abundance reached 6.6 ± 0.6 and 6.4 ± 0.6 × 106 cells ml^−1^ (mean ± standard deviations, n=3), respectively. This was nearly a doubling compared to the control cultures where 3.8 ± 0.7 × 106 cells ml^−1^ were recorded. Bacterial abundance in the *Brevundimonas* BAL450 cultures spiked with exponential phase DOM increased to 3.7 ± 0.4 × 106 cells ml^−1^, and further increased to 4.6 ± 0.4 × 106 cells ml^−1^ with stationary phase DOM. Control culture cell abundance was 1.7 ± 0.4 × 106 cells ml^−1^.

*Brief description of the Polaribacter sp. BAL334 and the Brevundimonas sp. BAL450 genomes Polaribacter* sp. BAL334 (Bacteroidetes) and *Brevundimonas* sp. BAL450 (Alphaproteobacteria) have similar genome sizes at ~3.3 Gb and ~3.2 Gb, respectively. The *Polaribacter* sp. BAL334 genome contains 2880 putative open reading frames (ORFs), of which 1021 (35.5%) have a SEED annotation, whereas 757 (26.3%) have a functional annotation but were not represented in SEED. The remaining 1102 ORFs (38.2%) were annotated as hypothetical proteins. The *Brevundimonas* sp. BAL450 genome encodes 3001 ORFs. Of these, 1269 (42.3%) have a SEED annotation, and 732 (24.4%) were not included in SEED but were functionally annotated; the remaining 1000 ORFs (33.3%) were annotated as hypothetical proteins.

As expected due to their role in central metabolism, dominant SEED categories (the highest level in the SEED hierarchy) in the genomes of both isolates were *Amino Acids and Derivatives* (up to 300 genes), *Carbohydrates*, *Protein Metabolism,* and the category *Cofactors, Vitamins, Prosthetic groups, Pigments* (**Fig. 2A**). Although the genome sizes of the two bacteria was comparable, *Polaribacter* BAL334 had a higher number of genomically encoded genes devoted to the categories *Carbohydrates* (222 genes, compared to 191 in *Brevundimonas* BAL450) and *Sulfur metabolism* (37 versus 18 genes) and *Photosynthesis*; the latter reflecting that only *Polaribacter* BAL334 has the proteorhodopsin gene encoded in its genome. The category *Motility and Chemotaxis* was only found in *Brevundimonas* BAL450 (84 genes), reflecting the lack of flagellar motility in *Polaribacter* BAL334 (**Fig. 2A**).

**Figure 2.**
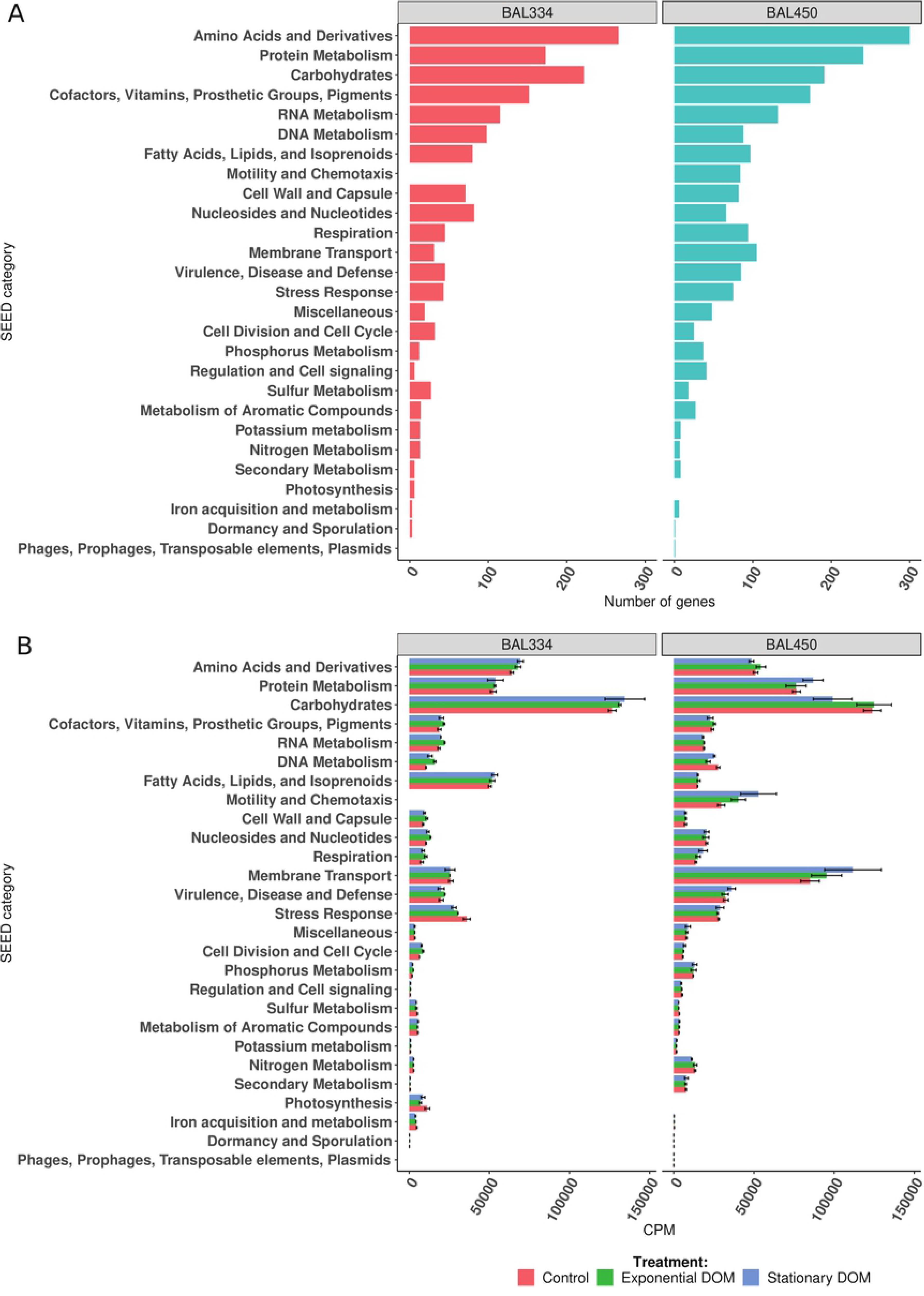
Comparison of number of genes in genomes and the relative expression levels in the two studied bacteria exposed to distinct DOM. A) Number of genomically encoded genes in top level SEED categories. B) Relative gene expression responses to enrichment with DOM collected from axenic exponential and stationary phase dinoflagellate cultures, and to control enrichments with only dinoflagellate L1 medium. “BAL334” and “BAL450” refers to *Polaribacter* BAL334 and *Brevundimonas* BAL450, respectively. Error bars in 2B indicate standard deviations for triplicates per treatment.

### Messenger RNA sequencing outcome

Sequenced bacterial mRNAs sampled 1 h after addition to phytoplankton DOC treatments mapped to 2742 ORFs in *Polaribacter* BAL334 and 2984 ORFs in *Brevundimonas* BAL450, representing 95% and 99% of putative protein coding genes in the genomes of the two isolates, respectively. The SEED annotated genes (with full functional hierarchies allowing higher level metabolic analyses; 995 and 1228 respectively in *Polaribacter* BAL334 and *Brevundimonas* BAL450), attracted 36-38% of the mRNA reads in *Polaribacter* BAL334 and 56-58% of the reads in *Brevundimonas* BAL450. The higher SEED annotation levels in the transcriptome of *Brevundimonas* BAL450 most likely reflects that knowledge of Proteobacteria genetics is generally higher than for Bacteroidetes.

### Overview of transcriptional differences between the two bacteria

Upon comparison of the two isolates, it was interesting to note that in the category *Membrane Transport*, both the number of genes in the genome (**Fig. 2A)** and the relative expression level (**Fig. 2B**) were threefold higher in *Brevundimonas* BAL450 than in *Polaribacter* BAL334 (105 genes versus 31 expressed genes at ~97,000 versus ~26,000 CPM). *Brevundimonas* BAL450 also had higher expression than *Polaribacter* BAL334 in the two categories *Nitrogen Metabolism* and *Phosphorus Metabolism* (**Fig. 2B**). Despite both genomes encoding around 90 genes in the category *Fatty Acids, Lipids, and Isoprenoids* (**Fig. 2A**), expression in this category was approximately 3-fold higher in *Polaribacter* BAL334 than in *Brevundimonas* BAL450 (~52,000 versus ~15,000 CPM) (**Fig. 2B**).

The major differences between the isolates in expression levels of genes in the SEED categories *Membrane Transport* and *Fatty Acids, Lipids, and Isoprenoids* led us to do a closer inspection of the genes involved. In *Membrane Transport*, we identified many more expressed genes in *Brevundimonas* BAL450 (64 paralog groups) than in *Polaribacter* BAL334 (18 paralog groups). Moreover, these genes were distributed in a broader variety of transporter subsystems (the second level in the SEED hierarchy; roughly corresponding to “pathways” in other annotation databases) in *Brevundimonas* BAL450 (**Fig. 3**). In both isolates, transporter expression was dominated by the *Ton and Tol transport* subsystem, both in relative expression levels and the number of expressed genes. *TRAP transporters* and *Uni-, Sym- and Antiporters* were well represented in *Brevundimonas* BAL450 but were absent from *Polaribacter* BAL334 (**Fig. 3**).

**Figure 3.**
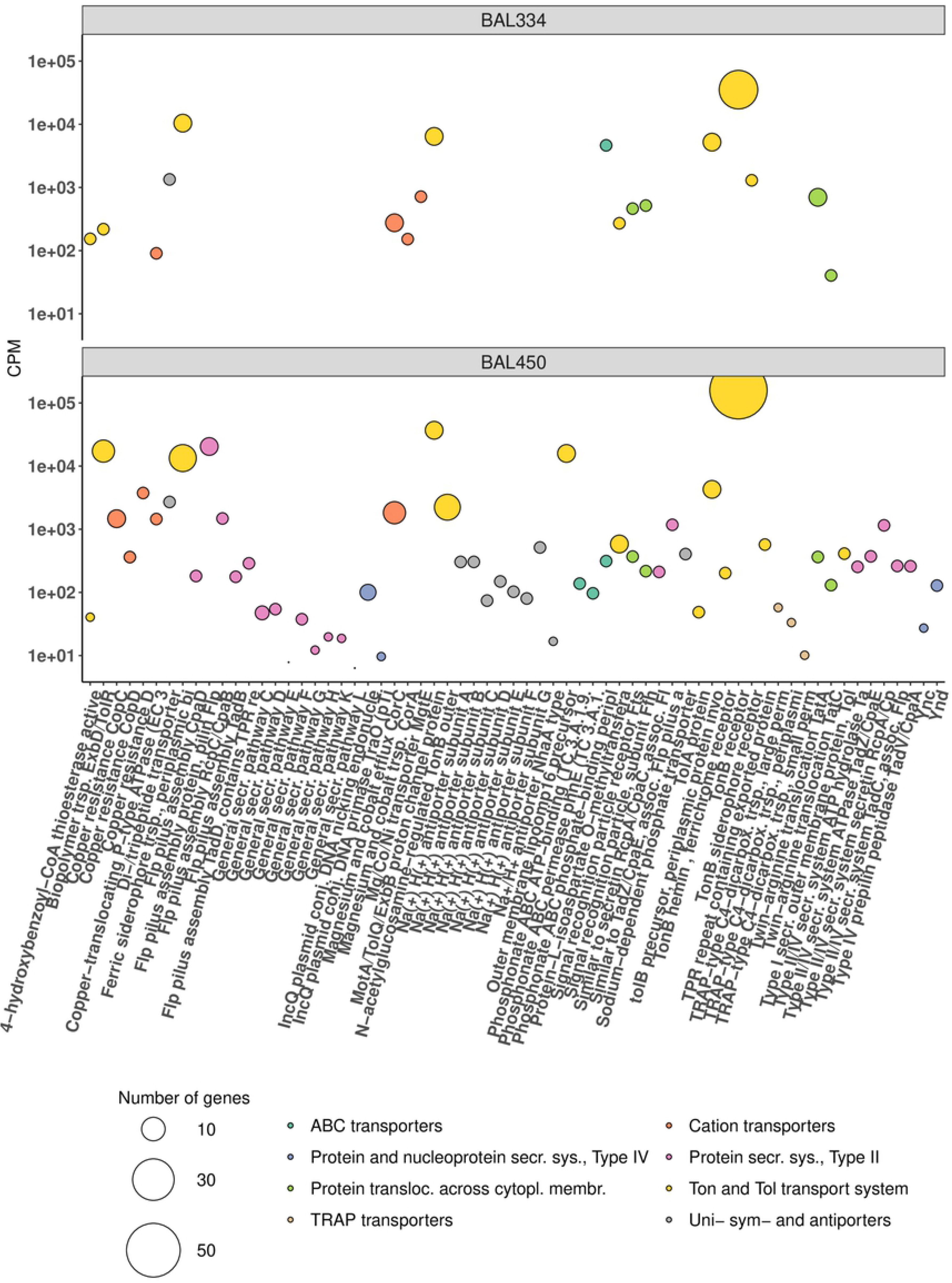
Comparison of relative expression levels of genes in the SEED category *Membrane Transport* in the two studied bacteria. Colors of circles denote transporter types. The size of circles represents the number of paralogs (defined as ORFs with the same name) in each of the genomes of the two bacteria. Y-axes shows the expression in counts per million (CPM) for the DOM treatments and the controls.

Regarding the category *Fatty Acids, Lipids, and Isoprenoids*, the higher expression in *Polaribacter* BAL334 compared to *Brevundimonas* BAL450 was primarily due to differences in *Fatty Acid metabolism* and *Polyhydroxybutyrate metabolism* (PHB) subsystems (**Fig. 4**). Both of the subsystems contained few expressed genes (five in both subsystems in *Polaribacter* BAL334; 11 and 15, respectively, in *Brevundimonas* BAL450; **Fig. 4**). In fact, two of the three most abundantly expressed genes in this category (*3-hydroxyacyl-CoA dehydrogenase* and *3-ketoacyl-CoA thiolase*) are shared between the subsystems, while the third (*Acetyl-CoA acetyltransferase*) occurred only in PHB. Still, we observed large differences between the isolates in the expression of these genes, with higher levels in *Polaribacter* BAL334 (**Table S1**). Moreover, *Polaribacter* BAL334 had substantially higher levels of expression of genes involved in isoprenoid synthesis (**Fig. 4**), in line with isoprenoids being precursors for carotenoids, which are likely responsible for the vividly orange color of *Polaribacter* BAL334 colonies.

**Figure 4.**
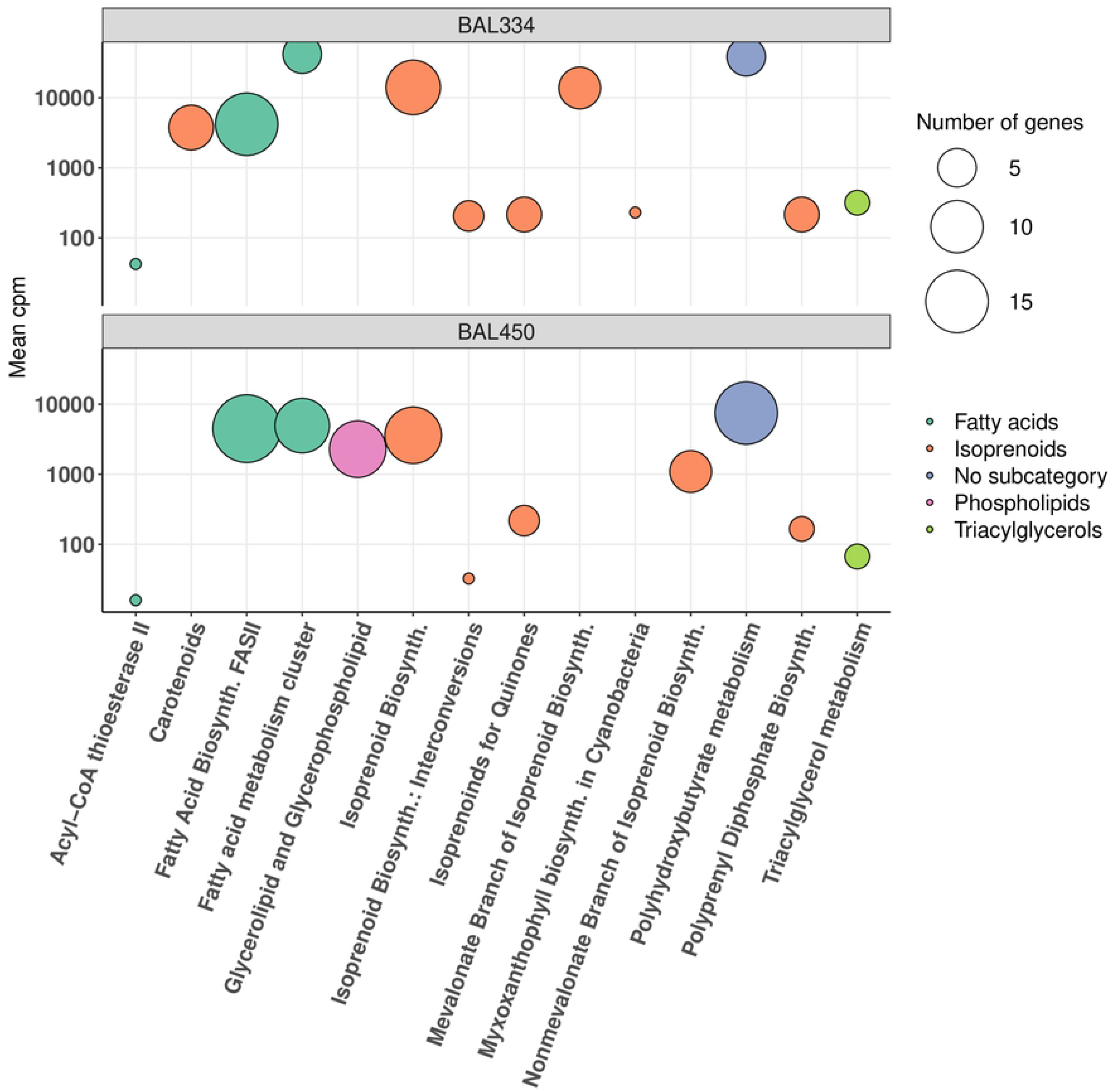
Analysis of gene expression in SEED subsystems of the *Fatty Acid, Lipids and Isoprenoids* category. Colors of circles denote SEED subcategories. The size of symbols represents the number of genes expressed in each subsystem. Y-axes shows the expression in counts per million (CPM) for the two DOM treatments and the controls.

### Significantly differentially expressed genes

To determine which expressed genes were significantly more (denoted “up”) or less (denoted “down”) abundant in the transcriptomes of the DOM-enriched samples than in controls, we performed a statistical analysis using EdgeR (29). The analysis was partitioned into four contrasts, considering each of the bacterial isolates and each of the DOM pools from the dinoflagellate relative to the control samples that did not receive any DOM. *Polaribacter* BAL334 enriched with DOM from exponential phase dinoflagellates compared to the control (hereafter, BAL334Exp-Con contrast) contained by far the highest number of differentially expressed genes (total of 128 genes; 106 up and 22 down; *false discovery rate <5%*) (**Fig. 5**). The treatment with DOM from stationary phase dinoflagellates (hereafter, *Polaribacter* BAL334Sta-Con) resulted in many fewer differentially expressed genes (24 genes up and two down). In *Brevundimonas* BAL450, on the other hand, it was the DOM from stationary phase dinoflagellates (hereafter, *Brevundimonas* BAL450Sta-Con) that induced more differentially expressed genes (23 genes up and ten down) than the exponential phase DOM (hereafter, *Brevundimonas* BAL450Exp-Con): four genes up and two down (**Fig. 5**). None of the significantly differentially expressed genes were shared between *Polaribacter* BAL334 and *Brevundimonas* BAL450 (**Table S2**). In both bacterial isolates the majority of expressed genes were not included in any SEED category (denoted “Not in SEED” in **Fig. 5**).

**Figure 5.**
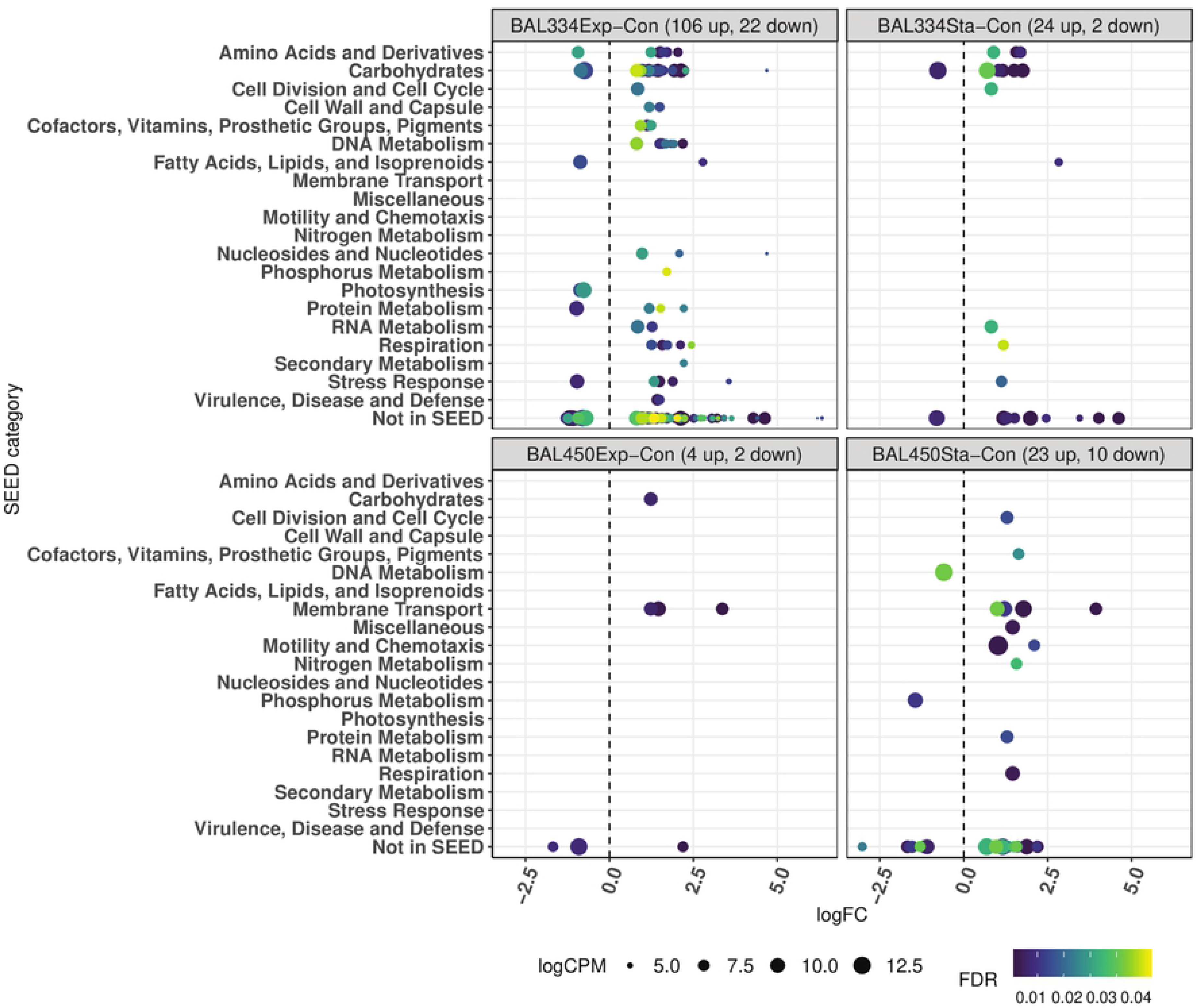
Influence of DOM from different growth phases of the dinoflagellate *Prorocentrum minimum* on statistically significant differences in gene expression between the two marine bacteria. Transcripts were defined as statistically significantly differentially abundant based on EdgeR analyses with an FDR ≤ 0.05. Each circle represents one gene and the circle size shows the calculated expression in logCPM; note that a gene can occur in more than one SEED category. Genes with significantly lower expression in controls compared to treatments are indicated with negative logarithmic fold change (logFC) values. BAL334Exp-Con and BAL334Sta-Con refers to *Polaribacter* BAL334 with DOM from exponential (Exp) and stationary (Sta) phase DOM compared to controls (Con). BAL450Exp-Con and BAL450Sta-Con refers to *Brevundimonas* BAL450 with DOM from exponential (Exp) and stationary (Sta) phase DOM compared to controls (Con).

Out of the 128 differentially expressed genes, 104 genes were unique to the *Polaribacter* BAL334 exponential phase. Twelve SEED subsystems had at least two differentially expressed genes compared to the control (**Fig. 6; Table S2**). Of these, only the *Proteorhodopsin* subsystem (in SEED category *Photosynthesis*) had genes with relative expression levels that were significantly higher in the controls (seen as negative value of differentially expressed genes in **Fig. 6**), one proteorhodopsin gene and one phytoene dehydrogenase (the latter being involved in the synthesis of the rhodopsin chromophore retinal). Both were moderately abundant with logCPM values of 11.1 and 9.6, respectively (i.e. ~0.2% and 0.08% of total transcripts).

**Figure 6.**
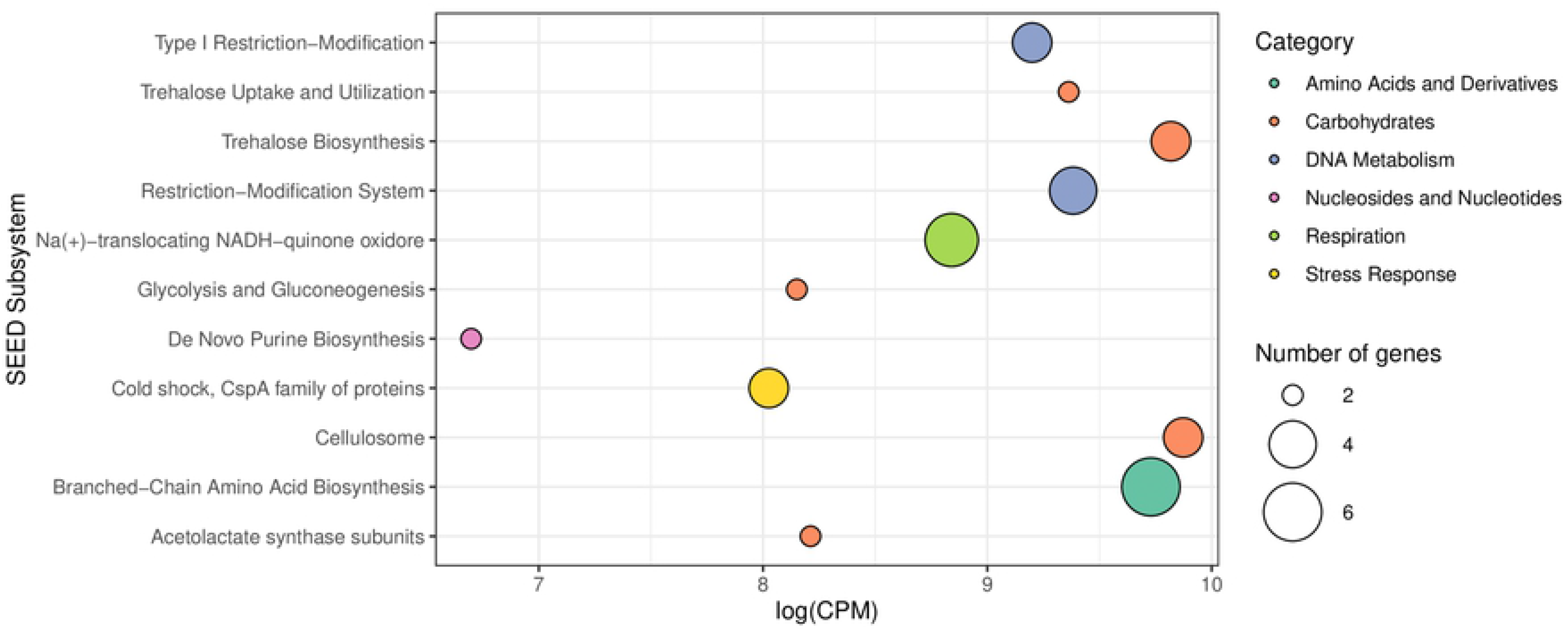
Subsystems in *Polaribacter* BAL334Exp-Con containing at least two significantly differentially expressed genes. SEED subsystem is shown on Y-axis and SEED category is shown in each plot title. The X-axis shows the number of genes whose expression is significantly more (positive value) or less (negative value) abundant compared with controls. Circle size represents the sum of logCPM.

Furthermore, the *Polaribacter* BAL334Exp-Con contrast contained five subsystems in the *Carbohydrates* category with at least two genes that increased in expression in the treated samples, three of which were also highly expressed: *Trehalose Uptake and Utilization*, *Trehalose Biosynthesis*, and *Cellulosome* (**Fig. 6; Table S2**). Several of the more abundant genes in these subsystems encode enzymes involved in binding of starch (SusC; a component of the polysaccharide utilization loci [PULs] widespread in Bacteroidetes (30), and eight genes involved in starch degradation to trehalose/maltose and the potential modification of these sugars: for example, two alpha-amylase genes the two enzymes in maltose to glucose degradation (maltose/trehalose phosphorylase **(Fig. 6; Table S2)**. This suggests that starch released by the dinoflagellate was an important substrate fueling growth of *Polaribacter* BAL334.

Finally, the *Polaribacter* BAL334Exp-Con contrast included the *Amino Acids and Derivatives* category, the *Branched-Chain Amino Acid Biosynthesis* subsystem which contained six genes involved in isoleucine synthesis. In the *DNA Metabolism* category, three type I restriction-modification system genes were detected, shared by two subsystems plus a type III restriction-modification system gene found only in the *Restriction-Modification System* subsystem (**Fig. 6; Table S2**). Only two significantly differentially abundant genes were unique to the contrast *Polaribacter* BAL334Sta-Con, encoding 2-isopropylmalate synthase (*Amino Acids and Derivatives*) and a SusC paralog, the outer membrane protein involved in starch binding (*Carbohydrates*).

Among the 32 significant genes that were unique in the *Brevundimonas* BAL450Sta-Con contrast, it is interesting to note the *Ferric siderophore transport system, periplasmic binding protein TonB* (**Table S2**). The Ton and Tol transport system (*Membrane Transport*) was the only subsystem in *Brevundimonas* BAL450 that contained at least two differentially expressed genes **(Fig. 6; Table S2)**. Two of the three genes were annotated as *TonB-dependent receptors*, and were shared between the exponential and stationary phase DOM contrasts with controls. In contrast, the *N-acetylglucosamine-regulated TonB-dependent outer membrane receptor* was unique to the BAL450Exp-Con contrast **(Fig. 6; Table S2)**.

## Discussion

We investigated how representatives of two major taxa in the marine environment – *Flavobacteria* and *Alphaproteobacteria* – react to DOM produced by dinoflagellates. These reactions partly reflected differences in genomically encoded functional capacity, but also that each of the isolates changed their allocation of relative transcriptional investment in certain metabolic functions. This change in transcription coincided with roughly a doubling in abundance following a single hour of DOM exposure. In agreement with the large phylogenetic distance between the two isolates, there were striking differences in the gene expression responses to DOM between the two isolates. These differences were noted even at the top level of the SEED classification system, with for example the ~3-4-fold higher relative expression of *Membrane Transport* in *Brevundimonas* BAL450, and the ~3-fold higher relative expression of *Fatty Acids, Lipids and Isoprenoids* in *Polaribacter* BAL334 compared to *Brevundimonas* BAL450. This suggests pronounced differences in resource utilization between these marine bacteria, indicating a potential for resource partitioning at the level of major metabolic categories.

As deduced from the number of genes that differed significantly in expression, we found that *Polaribacter* BAL334 (total 154 genes; 5.6% of genome) showed a much stronger response to DOM enrichment than *Brevundimonas* BAL450 (total 39 genes; 1.3% of genome). The responsiveness of *Polaribacter* BAL334 is in line with findings from the marine environment that flavobacteria have a pronounced ability to utilize organic matter produced during periods of phytoplankton blooms (31, 32). Moreover, it is noteworthy that the majority of significantly differentially expressed genes in *Polaribacter* BAL334 were observed in treatments with DOM from exponential phase dinoflagellates, whereas for *Brevundimonas* BAL450 most significantly expressed genes were found with stationary phase DOM. The latter observation was consistent with the observation that BAL450 reached higher bacterial abundances when incubated with stationary phase DOM. These findings imply that bacterial populations have diverged in their adaptation to utilize DOM produced and released by dinoflagellates during active growth compared to stationary phase. Our findings are encouraging for the future exploration of ecologically relevant patterns of how different bacterial taxa respond to and/or transform DOM produced by different phytoplankton species, and in relation to the physiological status of the phytoplankton as it differs across bloom development phases.

There were pronounced differences in which transporter genes the two bacteria expressed. For example, *Brevundimonas* BAL450 uniquely expressed a number of *Na+ H+ antiporters* (involved in pH and/or salinity adaptation (33)) and secretion system transporters (involved in adhesion, e.g. to algal cells (34)). Curiously, one of the most highly expressed genes in *Polaribacter* BAL334 was a transporter for phosphonate (also expressed in *Brevundimonas* BAL450 but at lower levels).

Phosphonate is an organic form of phosphorus which can be used as a sole source of phosphorus by some microorganisms, allowing them higher fitness under phosphorus limiting conditions (35–37). It is estimated that phosphonate constitutes a large fraction (5-25%) of the dissolved organic phosphorus (DOP) in the oceans (38). During phosphate depletion in phytoplankton blooms, ABC-type phosphonate transporters proteins typically increase in abundance in some bacterial taxa (32). Genes involved in phosphonate utilization are thus candidates to act as sensors for phosphate status in marine environments, complementing genes involved in phosphate utilization (e.g. phosphate membrane transporters and alkaline phosphatase) (39).

In the two model bacteria studied here, we found that the expression of *Ton and Tol transport systems* were dominant in both transcript abundance and in number of expressed genes. This class of transporters is found in the outer membrane of gram-negative bacteria and is involved in the uptake of a broad set of macromolecules, such as siderophores for iron, vitamin B_12_, nickel complexes and poly- or oligomeric carbohydrates (40). Since DOM produced by phytoplankton can be rich in polysaccharides amenable to utilization by bacteria (41), expression of transporters for this type of compounds can be expected. It is particularly intriguing that we found so many different genes involved in the Ton and Tol systems expressed, as this is consistent with the uptake of not just a few preferred molecules, but the simultaneous uptake of a wide array of compounds exuded by phytoplankton. The characterization of sets of transporters in greater detail thus has the potential to provide deeper understanding of bacterial DOM metabolism along the progression of phytoplankton blooms.

Interestingly, *Polaribacter* BAL334 also had higher relative expression of genes involved in the subsystems *Fatty acid metabolism cluster* and *Polyhydroxybutyrate metabolism* (**Fig. 3**; **Table S1**). PHB is produced by diverse bacteria in response to physiological stress or carbon excess (42). The carbon stored in PHB can be used later as an energy source or as anabolic building blocks in times of low availability of DOM (42). Interestingly, another flavobacterium, *Dokdonia* sp. MED134, has earlier been seen to express genes for a different carbon storage molecule – glycogen – under conditions where two strains of proteobacteria expressed genes for PHB synthesis (43). *Polaribacter* BAL334 encodes both pathways and the carbon storage strategy hence appears to not only reflect phylogenetic relatedness but also temporary ecological factors such as the composition of available substrates.

Even at the highest level of the SEED hierarchy, two categories stood out being exclusively expressed by just one of the two isolates: *Motility and Chemotaxis* in *Brevundimonas* BAL450 and *Photosynthesis* in *Polaribacter* BAL334 (**Fig. 2B**). Genome analysis showed that *Polaribacter* BAL334 lacks the full complement of flagellar motility (it uses gliding motility for movement) (44). In contrast, the flagellar motility system is present in *Brevundimonas* BAL450 where it was highly expressed. At the top level SEED (**Fig. 2B**), motility and chemotaxis gene expression was particularly high in the treatments with dinoflagellate DOM as compared to controls. This could potentially relate to the cells sensing increased nutrient availability in the DOM treatments or that the DOM provided energy that fueled increased swimming (45). The *Photosynthesis* genes expressed by *Polaribacter* BAL334 are those encoding the energy-generating proteorhodopsin photosystem, which *Brevundimonas* BAL450 lacks. Strikingly, the two differentially abundant genes in proteorhodopsin synthesis were expressed at higher relative values in the controls than in the treatments with DOM. Proteorhodopsin is known to help bacterial cells to survive during starvation (46) or even contribute to growth at low DOC availability (47). This suggests that *Polaribacter* BAL334 used the proteorhodopsin for surviving in the no-substrate controls, and when provided with DOM in the treated samples preferentially utilized DOM rather than photoheterotrophy for its energy demand.

The relative expression responses we observed between bacterial species, and between DOM from two different dinoflagellate growth phases, helped identify genes potentially involved in shaping the ecology of heterotrophic marine microbes. Our findings emphasize the potential usefulness of experimental approaches for identifying indicator genes for different environmental conditions that are informative of mechanisms underlying important dynamics of carbon and nutrient fluxes in marine ecosystems. Our findings have implications for metatranscriptome analysis, since sequences taken from a community of phylogenetically diverse populations will likely blur signals of biogeochemical relevance because of differences in functional capacity and lifestyles between species. Separation of taxa based on taxonomic annotation before analysis of differentially abundant genes has been proposed to resolve this issue (48). Attaining sufficient precision in the identification of species – for example through the use of metagenomic assembled genomes (MAGs) (49, 50) – would allow the use of the genetic responses of particular species of marine bacteria sampled in natural environments as “living sensors”.

## Acknowledgments

We thank Sabina Arnautovic for skillful technical assistance. Computations were performed on resources provided by the Swedish National Infrastructure for Computing (SNIC) through the Uppsala Multidisciplinary Center for Advanced Computational Science (UPPMAX) under project SNIC 2017/7-419. DNA sequencing was conducted at the Swedish National Genomics Infrastructure (NGI) at Science for Life Laboratory (SciLifeLab) in Stockholm.

## Author contributions

JP, MAM and CMGO conceived the study. CMGO and CK performed the experiment. Laboratory work was done by CMGO and CK with help from JET. DL and CMGO did the bioinformatics. CMGO, DL, MAM and JP wrote the article.

## Supplementary table legends

**Table S1.** Expression of genes in the SEED category *Fatty Acids, Lipids, and Isoprenoids* in *Polaribacter* BAL334 and *Brevundimonas* BAL450. Both the SEED subcategory and subsystem are shown together with the expression abundance of the gene in counts per million (CPM) and standard deviation (CPM) in treatments and control. Note that a gene can occur in more than one SEED category.

**Table S2.** Significantly differentially abundant genes in *Polaribacter* BAL334 and *Brevundimonas* BAL450 with SEED categories and SEED subsystems. Expressed genes were determined to be significantly more (denoted “up”) or less (denoted “down”) abundant in the transcriptomes of the DOM-enriched samples compared to controls, using EdgeR statistical analysis. Contrast indicates whether the gene occurs only in DOM from exponential phase (i.e. Exp-Con) or only in the stationary phase (i.e. Sta-Con) or is in both (i.e. shared genes). Note that a gene can occur in more than one SEED category.

